# Functional Roles for RNA Ribosylation in Mammalian Cells

**DOI:** 10.1101/2025.07.18.665567

**Authors:** John D. Sears, Che-Kang Chang, Takayoshi Shirasaki, Rebekah Dickmander, Nicholas A. Saba, Wes Sanders, Mark T. Heise, Nathaniel J. Moorman

## Abstract

In mammalian cells, the addition of ADP-ribose to proteins and DNA plays well established roles in regulating cell function. Recently, RNA ribosylation was also found in mammalian cells under conditions of cell stress, though the functional consequences remain unclear. Here we find that infection with chikungunya virus, a positive strand RNA virus that causes frequent widespread epidemics, increases overall levels of RNA ribosylation in human fibroblasts. During infection, viral RNA is ribosylated by the PARP12 ribosyltransferase, which is counteracted by a virally-encoded. Increased viral RNA ribosylation resulted in decreased translation in cell-free systems and infected fibroblasts, and more rapid viral RNA decay. Further, ribosylated RNA potently induced the expression of antiviral host response genes. Together these data show the first functional consequences of RNA ribosylation in mammalian cells by showing that RNA ribosylation inhibits translation, decreases RNA stability, and creates a novel pathogen-associated molecular pattern (PAMP) that activates the host innate immune response. As macrodomains are present in multiple unrelated viruses, our data suggest RNA ribosylation is a novel component of cellular antiviral sensing pathway. These results also provide a starting point for defining functional roles for RNA ribosylation in other mammalian cell stress conditions beyond viral infection.

## INTRODUCTION

RNA modifications play critical roles in regulating RNA activity and metabolism at multiple levels. For example, the addition of the 5′ 7-methylguanosine cap and its subsequent methylation modulate RNA translation, while other modifications such as methylation and acetylation impact RNA stability, localization, and degradation (1–4). RNA modifications are especially important under conditions of cellular stress, such as viral infection, where they help integrate environmental signals and determine RNA fate (5, 6). Understanding the functional consequences of RNA modifications is therefore critical for uncovering how cells adapt to stress and dynamically reprogram gene expression to promote cell survival.

ADP ribosylation, the covalent attachment of adenosine diphosphate ribose (ADPr) to substrates, has long been recognized as a conserved mechanism for responding to stress across diverse biological systems. Both proteins and nucleic acids can be ADP-ribosylated, and modification of both regulates cellular stress responses (7). For example, in bacteria and plants, ribosylation of exposed DNA ends is recognized as a damage signal and helps recruit DNA repair proteins. Protein ribosylation of recruited factors further facilitates chromatin remodeling and promotes error-free DNA repair in bacterial, plant, and mammalian cells (7). During viral infection, protein ADP ribosylation also plays key roles by attaching ADPr to essential viral replication proteins to inhibit their function (8, 9). Additionally, ribosylation of host innate immune effectors can enhance their antiviral activity (10). These examples underscore the evolutionary conservation and functional importance of ADP ribosylation in regulating cellular responses to diverse stressors.

In addition to DNA and protein substrates, RNA has recently been identified as a target of ADP ribosylation, though its function in mammalian cells is unknown. In prokaryotes, RNA ribosylation serves as an antiviral defense mechanism against bacteriophage infection. For example, bacterial cells upregulate the ADP-ribosyltransferase CmdT upon bacteriophage infection, which modifies the 5′ end of phage RNA to block its translation and thereby inhibit viral replication (11). In mammalian systems, cellular stressors that inhibit translation such as nutrient deprivation or the accumulation of unfolded proteins also induce RNA ribosylation (12, 13). In cell-free systems RNA modified with ADPr on the 5’ terminus in place of the 7-methylguanosine mRNA cap reduces translational efficiency and protects from degradation by the XRN-1 exoribonuclease (12). These observations, along with its known antiviral role in bacteria, suggest that RNA ribosylation may similarly contribute to stress adaptation in mammalian cells. However, to date, no physiological role for RNA ribosylation has been reported in any mammalian system.

RNA ribosylation is catalyzed by RNA ADP-ribosyltransferases, which utilize NAD⁺ as a substrate to covalently add ADPr to specific sites on RNA molecules (12, 14). ADPr can be attached to various locations on RNA, including 5′ and 3′ terminal phosphates (13), the exocyclic amine of guanosine bases (15), and the 2′ hydroxyl group of ribose (16). ADP ribosyltransferases are conserved across all domains of life, supporting the notion that RNA ribosylation is a fundamental and ancient regulatory mechanism (17–19). In cell-free systems, several mammalian ADPr ribosyltransferases, including multiple members of the poly-ADPr polymerase (PARP) family such as PARP10, 11, 12, 15, and TRPT1, can ribosylate RNA (12). In mammalian cells these ADPr ribosyltransferases are required for RNA ribosylation in response to cellular stress. Although the physiological consequences of this modification remain unknown in mammals, its conservation and induction under stress suggest important regulatory roles, possibly analogous to its functions in prokaryotic systems.

Specialized ribosylhydrolases remove ADPr moieties from RNA by catalyzing the hydrolysis of the glycosidic or diphosphate bond. The NADAR family found in bacteria, archaea, viruses, and certain eukaryotes specifically removes ADPr from guanosine bases (16), however NADAR orthologs are absent in mammals (20). Both ARH and macrodomain ribosylhydrolases are expressed in mammals and can remove ADPr from 2′ hydroxyl groups and terminal phosphates on RNA in cell-free systems (16, 21). Several viruses, including coronaviruses (e.g. SARS-CoV-2) and alphaviruses (e.g. Chikungunya virus), encode macrodomains that efficiently remove ADPr from RNA in cell-free systems (16, 21), suggesting that viral ADPr ribosylhydrolases may suppress RNA ribosylation during infection.

Here we explore functional roles for RNA ribosylation in mammalian cells, using Chikungunya virus (CHIKV) infection of human fibroblasts as a model system. Our results show that RNA ribosylation serves as a novel cellular defense during viral infection by inhibiting viral RNA translation, inducing viral RNA decay, and stimulating the innate antiviral response to infection. Our data provide new insights into how RNA ribosylation integrates with known cellular signaling networks such as the type I interferon pathway, suggesting a central role for RNA ribosylation in the response to infection. More broadly, our results provide a framework for investigating how RNA ribosylation may more generally regulate cell biology during periods of cell stress.

## MATERIAL AND METHODS

### Cell Culture and Virus Production

Human MRC-5 hTert fibroblasts were grown in Dulbecco’s Modified Eagle Medium (DMEM, Gibco) supplemented with 10% fetal bovine serum (FBS, Gibco) and 1% L-glutamine (Gibco). BHK-21 cells were grown in MEM-alpha media (Gibco) supplemented with 10% FBS, 1% L-glutamine, and 5% tryptose phosphate broth. All cells were cultured 37 °C in 5% CO_2_. Viruses were produced by electroporating BHK-21 cells with in vitro transcribed, capped RNA from an infectious clone of CHIKV (181/25). Briefly, an infectious clone of 181/25 was linearized with NotI-HF (NEB), purified, and used as a template for in vitro transcription using mMessage Machine Kit (Invitrogen) according to the manufacturer’s directions. In vitro transcribed RNA was precipitated with LiCl, and then 10µg of RNA was electroporated into 4x10^7 BHK-21 cells. Virus was purified from cell supernatants 24 hours after electroporation by centrifugation through a 20% sucrose cushion. The resulting pellet was resuspended in PBS supplemented with 1% FBS and stored at -80°C until use.

### Western Blotting

Protein was extracted from cells using RIPA buffer (50 mM Tris-HCl [pH 7.4], 150 mM NaCl, 1 mM EDTA, 1% NP-40, 1% sodium deoxycholate) supplemented with protease inhibitor cocktail (Roche cOmplete EDTA-free protease inhibitor cocktail) as previously described (22). Protein concentration was determined using a Bradford assay (BioRad). 30 µg of protein was resolved on 10% SDS-PAGE gels and then transferred to a PVDF membrane. Membranes were blocked with 5% non-fat dairy milk in TBST (20 mM Tris-HCl [pH 6.8], 140 mM NaCl, 0.1% Tween 20) for 1 hour prior to incubation with primary antibody overnight at 4°C in TBST + 5% BSA. Blots were washed with TBST prior to incubation with the appropriate secondary antibody conjugated to horseradish peroxidase (Advansta) in TBTS + 5% BSA and visualized by chemiluminescence. Antibodies used in this studies include: Poly/Mono ADPr (Cell Signaling Technology; #E6F6), CHIKV nsP2 (Invitrogen; #HL1432), Tubulin (Cell Signaling Technology; #2144), PARP10 (Bethyl; #A300-665A), PARP11 (Proteintech; #16692-1-AP), PARP12 (Invitrogen; #PA5-89749), PARP15 (Proteintech; #18126-1-AP), CHIKV Capsid (Invitrogen; #PA5-143450), GAPDH (Cell Signaling Technology; #14C10), Anti-Rabbit HRP IgG (Seracare Life Sciences #54500010), Anti-Mouse HRP IgG (Seracare Life Sciences #54500011).

### Dot blot assay

Ribosylated RNA was detected following previously described protocols (23). In brief, RNA was extracted using RNAeasy Kits (Qiagen) according to the manufacturer’s directions. RNA was dotted onto nylon membrane (Invitrogen) and allowed to dry for 15 minutes prior to UV cross-linking using the auto-cross link function of the Stratalinker (Stratagene). Membranes were blocked with 5% milk in PBST prior to incubation overnight with Poly/Mono-ADPr antibody in PBST supplemented with 5% BSA. Membranes were washed with PBST and probed with secondary antibody in 3% BSA prior to visualization via chemiluminescence. For experiments to evaluate ADPr antibody specificity, RNA was extracted as above from human fibroblasts at 18 hours after infection with CHIKV (181/25). 1µg of RNA was treated with either 1 unit Proteinase K (NEB), 1µg RNAse A (Invitrogen), or 2 units of DNAse (Invitrogen) for 2 hours at 37°C, and then re-purified. An equal volume of each sample was dotted onto nylon membrane to detect ribosylation as above.

### Infectious Virus Quantification

The specific infectivity of viral RNA was measured as previously described (24). Briefly, BHK-21 cells were electroporated with in vitro transcribed RNA as described above. Serial 10-fold dilutions of electroporated cells were plated onto monolayers of Vero81 cells and then overlayed with 2x α-MEM (Gibco) containing 2.5% carboxmethylcellulose. Seventy-two hours after plating, cells were fixed with 2% paraformaldehyde and stained with crystal violet to visualize plaques. Plaque assays were used to quantify levels of cell-free infectious virus, as previously described (24). Briefly, serial 10-fold dilutions of virus stocks were plated in Vero81 cells. After adsorption for one hour at 37°C, the inoculum was removed and replaced with 2x α-MEM (Gibco) containing 2.5% carboxmethylcellulose, incubated for 72 hours and were fixed and stained as above.

### Quantification of Viral Genomes in Virus Stocks

Viral genomes were quantified by RT-qPCR as described previously (25). Briefly, RNA was extracted from 100uL of concentrated virus stocks using Trizol. Viral RNA was reverse transcribed and quantified by RT-qPCR using primers specific for nsP2 *(F 5’-TGCGTGAGACACACGTAGCCT-3’, R 5’-GGGCCTTCAAAAAGGCGCTGTCA-3’)*. The number of viral genomes was determined by the absolute quantification method by comparing the results to a standard curve generated using a range of viral genomes from 10^8 to 10^1 copies.

### RNA Immunoprecipitation

RNA was isolated from human MRC-5 fibroblasts at 18 hours after infection using an RNAeasy kit (Qiagen) according to the manufacturer’s directions. Mono/Poly-ADPr antibody (Cell Signaling Technologies; #E6F6a) or an IgG control was conjugated to protein A/G PLUS agarose beads (Santa Cruz). 5µg of RNA was incubated with the beads for 1 hour, after which the beads were washed with a series of buffers of increasing stringency to minimize non-specific binding, as before (26). RNA was extracted from the beads with Trizol, and the number of viral genomes quantified by RT-qPCR using the absolute quantification method and nsP2-specific primers noted above.

### RT-qPCR quantification of RNA expression

For RNA decay and ISG expression experiments RNA was extracted from MRC-5 fibroblasts using the RNAeasy kit (Qiagen) as described above. 1 µg of total RNA was reverse transcribed using the High Capacity cDNA Reverse Transcription Kit (Applied Biosystems #4368813) according to the manufacturer’s directions. Viral RNA was quantified via absolute quantification method using nsP2-specific primers as described above. Host genes were quantified using the ΔΔCT method normalized to actin. Primers sequences are listed below: nsP2 *(F 5’-TGCGTGAGACACACGTAGCCT-3’, R 5’-GGGCCTTCAAAAAGGCGCTGTCA-3’),* Capsid (F 5’-GACCGATCTTCGACAACAAG-3’, R 5’-CGATGTCTTTGTTCCAGGTC-3’), ISG15 (F 5’-TGGCGGGCAACGAATT-3’, R 5’-GGGTGATCTGCGCCTTCA-3’), IFIT1 (F 5’-GCCTTGCTGAAGTGTGGAGGAA-3’, R 5’-ATCCAGGCGATAGGCAGAGATC-3’), RIG-I (F 5’-CACCTCAGTTGCTGATGAAGGC-3’, R 5’- GTCAGAAGGAAGCATTGCTACC-3’), OAS1 (F 5’-AGGTGGTAAAGGGTGGCTCC-3’, R 5’-ACAACCAGGTCAGCGTCAGAT-3’).

### Rabbit Reticulocyte Lysate Translation Assay

50 ng of RNA was incubated at 65°C for 5 minutes prior to incubation on ice. The RNA was then added to nuclease treated rabbit reticulocyte lysates (Promega #L4960) supplemented with 50 mM KCl, 20 µM unlabelled amino acids, and 25 units RNAse inhibitor (Promega #N2111). Reactions were incubated at 30°C for 90 minutes and 5µL was removed at 0, 5, 15, 30, 45, 60, and 90 minutes and immediately treated with Nano-Glo reagent (Promega #N1110) and luciferase activity was quantified using a Glomax plate reader (Promega).

### XRN-1 Degradation Assay

Purified virion RNA or control mouse thymus RNA was treated with either 10U RppH (NEB, cat # M0356) or with buffer as negative control. The reaction was incubated at 37°C for 1 hour and then re-purified as described above. 2U XRN-1 (NEB; #M0338) or buffer was added to each sample and incubated at 37°C for 1 hour. Equal volumes of each reaction were analyzed by RT-PCR using the nsP2-specific primers described above and quantified by the absolute quantification mehod. The percentage of XRN-1 resistant RNA was calculated by dividing the number of viral genomes in XRN-1 treated samples by the number of viral genomes in buffer-treated samples and multiplying by 100.

### PARP siRNA Knockdown

MRC-5 hTert human fibroblasts were reverse transfected with SMARTPool ON-TARGETplus siRNAs (Dharmacon) specific for the indicated genes, non-targeting siRNAs, or a combination of all PARP siRNAs together as previously described (26). 48 hours after transfection, cells were infected at 48 hours after transfection and cells and culture supernatants were harvested at 24 hours after infection. Total RNA was extracted using the RNeasy Kit, and RT-qPCR was performed using nsP2 specific primers as above to measure viral RNA abundance. Knockdown efficiency and impact on viral replication were measured by Western blot and RT-qPCR.

## RESULTS

### Virus Infection Increases RNA Ribosylation

Recent studies have found that RNA can be modified with ADP ribose (ADPr RNA) in mammalian cells and that RNA ribosylation is induced under conditions of cell stress such as nutrient deprivation and heat shock (12). As viral infection also activates a cell stress response, we tested if infection with chikungunya virus (CHIKV), a positive sense RNA virus in the Togavirus family that causes widespread, explosive outbreaks and epidemics of severe acute and chronic arthralgia, induced RNA ribosylation in infected cells. Total RNA and protein were extracted from CHIKV infected cells or uninfected control cells and protein and RNA ADPr levels were measured by Western blot and dot blot (23), respectively. Infection led to a consistent increase in ribosylated protein and RNA levels compared to controls (Figure 1A, B). To confirm that the signal from the dot blot was specific for ribosylated RNA rather than contamination from ADPr modified DNA or protein, we treated total RNA from infected cells with RNAse A, DNase, or Proteinase K, and then re-purified the RNA and again measured ribosylated RNA levels. Only samples treated with RNAse A displayed a significant decrease in the ADPr RNA signal, demonstrating that the assay specifically measures ribosylated RNA rather than DNA or protein (Figure 1C). Together these data show that like other cell stressors, viral infection induces RNA ribosylation in mammalian cells.

**Figure 1.**
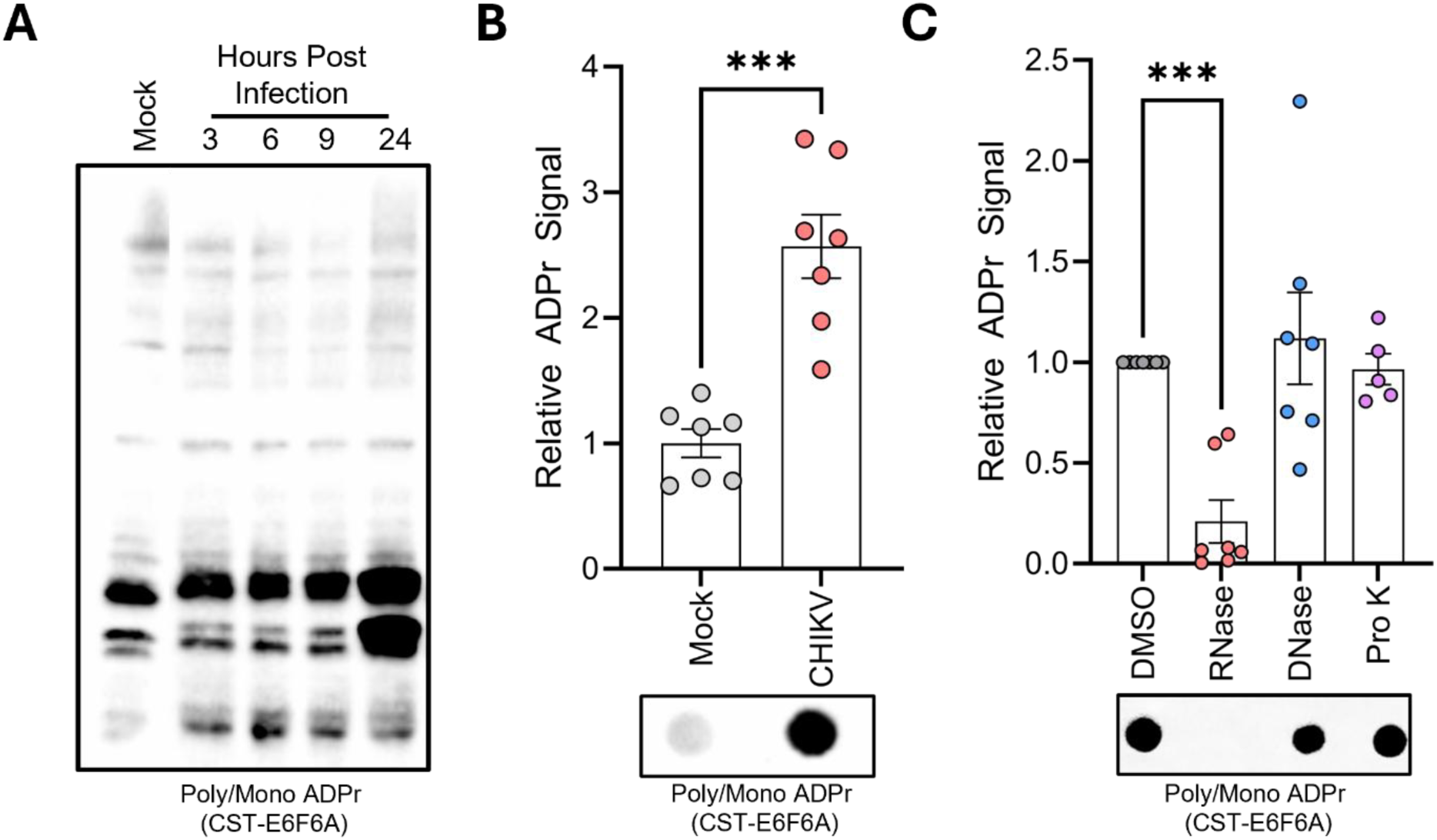
Chikungunya virus infection increases RNA ribosylation. (A) Human fibroblasts were infected with chikungunya virus (CHIKV) strain181/25 at an MOI=3 and harvested at the indicated times after infection. Protein ribosylation was determined by Western blot of infected cell extracts. (B) Total RNA was extracted from cells infected as above and harvested at 18 hours after infection. The levels of ribosylated RNA were determined by dot blot. (C) RNA from (B) was treated with DMSO, 1µg RNAse A, 2 units DNAse, or 1 unit proteinase K for 2 hours. ADP-ribose levels were measured by dot blot and quantified relative to DMSO treated samples (N=3, ***=p<0.001).

### The CHIKV nsP3 Macrodomain Reverses RNA Ribosylation in Mammalian Cells

The CHIKV nsP3 protein contains a macrodomain, an evolutionarily conserved ADPr ribosylhydrolase domain essential for efficient viral replication, that removes inactivating ADPr modifications from viral proteins (27–29). Together with the increase in ADPr RNA observed after infection, the presence of a viral macrodomain suggests that the CHIKV nsP3 macrodomain may limit ribosylated RNA accumulation during infection. In Sindbis Virus (SINV), a related alphavirus, macrodomain activity is essential for virus replication, however mutations that attenuate macrodomain activity are still viable. Specifically, an asparagine to alanine (N24A) or glycine to glutamic acid (G32E) mutation in the SINV macrodomain active site decreases ribosylhydrolase activity and limits virus replication (28). To determine if analogous mutations in CHIKV similarly impacted virus replication, we measured the replication of a set of mutations to the CHIKV nsP3 macrodomain compared to wild type virus. We determined that the nsP3 N24A mutation was the only mutation that displayed a significant decrease in specific infectivity and replication without signs of reversion and therefore used it for subsequent analyses (Supplemental Figure 1A-E). We first measured the impact of N24A on replication using a specific infectivity assay, which quantifies the number of productively infected cells after electroporation and provides a measure of whether a given mutation adversely affects the ability of the viral RNA to initiate infection (Figure 2A, Supplemental Figure 1C) and the yield of infectious virus 24 hours following electroporation (Figure 2B, Supplemental Figure 1D). As with SINV, the N24A mutation significantly decreased RNA specific infectivity and cell-free virus yield at 37°C. However, this defect could be rescued when transfected or infected cells were incubated at 30°C, demonstrating that the defect was temperature dependent (Figure 2A, B).

**Figure 2.**
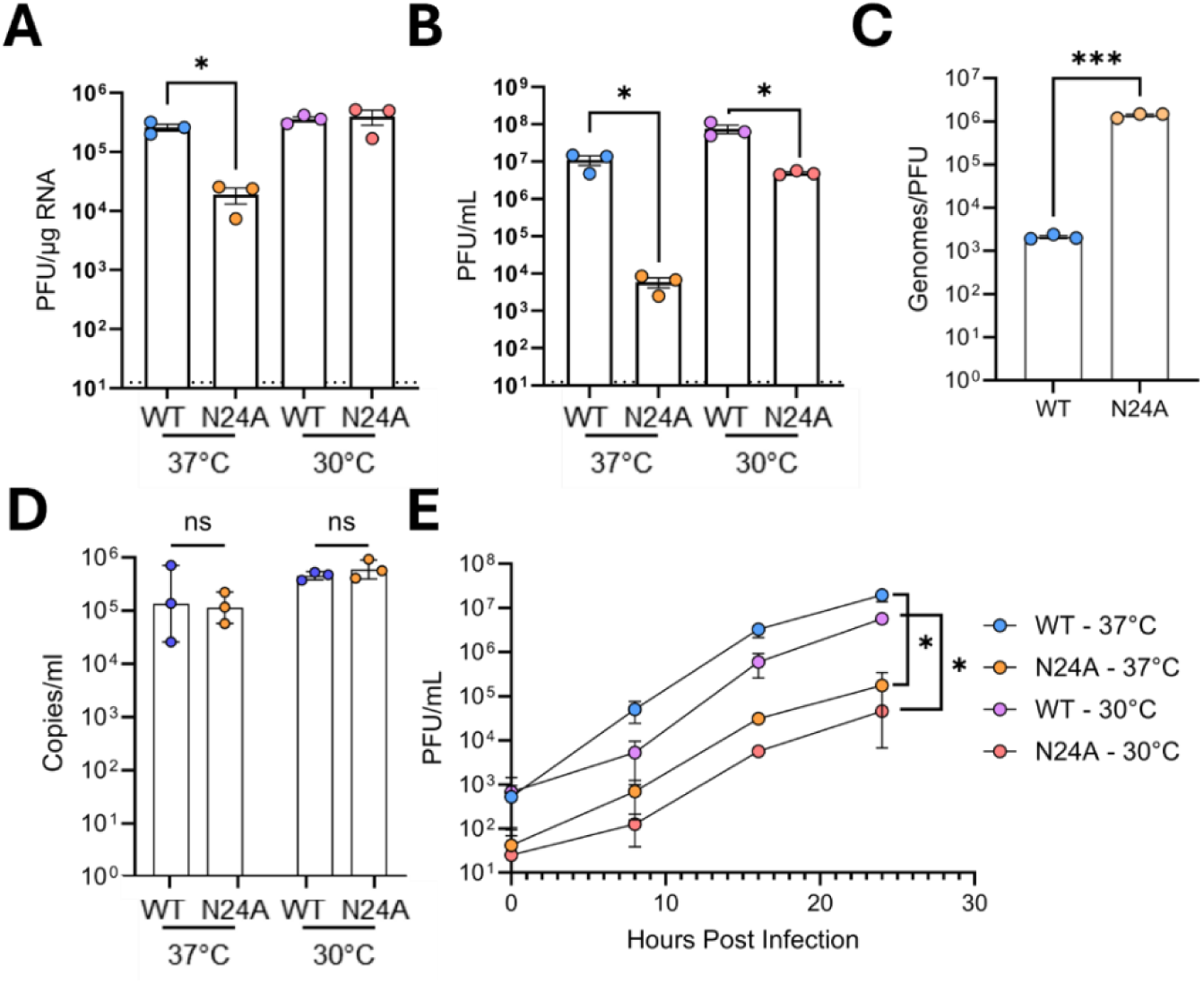
CHIKV nsP3 macrodomain is required for efficient virus replication. (A) BHK21 cells were electroporated with wild type (WT) or N24A RNA and the specific infectivity of WT and N24A was determined by plating electroporated BHK21 cells on a monolayer of Vero81 cells. (B) Supernatants were collected 24 hours after electroporation and titered on Vero81 cells at either 37°C or 30°C (C) The ratio viral genomes to plaque forming units (PFU) was determined by measuring genome copies from viral supernatants by RT-qPCR and comparing it to viral titers at 37°C (D) Human fibroblasts were infected with an equivalent number viral genomes of WT or N24A virus, which was equal to a multiplicity of infection of 3 PFU/cell for wild type virus. Viral RNA levels in infected cells was quantified by RT-qPCR at 1 hour after infection. (E) Human fibroblasts were infected with the equivalent number of genomes of WT and N24A as in D, and amount of infectious virus at the indicated times after infection was measured by plaque assay. (N=3, *=p<0.05, ***=p<0.001).

To determine if the different replication rates of N24A as compared to wild type virus were caused by an accumulation of defective genomes, we assessed the ratio of viral RNA to the number of plaque forming units in N24A or wild type virus stocks as measured by plaque assay. We observed a significant disconnect between the number of viral genomes in virus stocks when compared to the number of infectious particles. The N24A and wild type virus stocks containing similar numbers of viral genomes, but the N24A virus stocks had a significant defect in the number of infectious particles (Figure 2C). While this difference could be due to a defect in viral binding and entry, we found that when cells were infected with an equal number of N24A and wild type viral genomes, equivalent numbers of viral genomes were found inside the cell at 1 hour after infection, demonstrating equivalent binding and entry for both viruses (Figure 2D). As a further measure of virus replication, we infected cells with an equal number of N24A and wild type viral genomes analyzed viral replication kinetics by measuring the accumulation of infectious virus across a time course of the viral replicative cycle. As we found different specific infectivity phenotypes for the N24A mutant at different temperatures, we repeated the experiment at both 37° and 30°C. Wild type CHIKV replicated more efficiently at 37°C as compared to 30°C. As with the analogous SINV mutant, the CHIKV N24A mutant replicated less efficiently than wild type virus, and this defect was significant at both 37° and 30°C (Figure 2E). We conclude that, similar to SINV, nsP3 macrodomain activity is required for efficient CHIKV replication, and attenuating CHIKV nsP3 macrodomain activity results in inhibition of virus replication at a step after virus binding and entry.

The SINV nsP3 macrodomain removes inactivating ADPr from viral and cellular proteins to enhance virus replication, and mutating nsP3 at N24A significantly reduces its ribosylhydrolase activity (27). To confirm that the N24A mutation in CHIKV similarly decreased nsP3 macrodomain activity, we measured the accumulation of ribosylated proteins in human fibroblasts infected with wild type CHIKV or the N24A CHIKV mutant. As expected, increased levels of protein ribosylation were observed in N24A infected cells as compared to cells infected with wild type virus, confirming that the N24A mutation decreased ribosylhydrolase activity of the CHIKV nsP3 macrodomain (Figure 3A).

**Figure 3.**
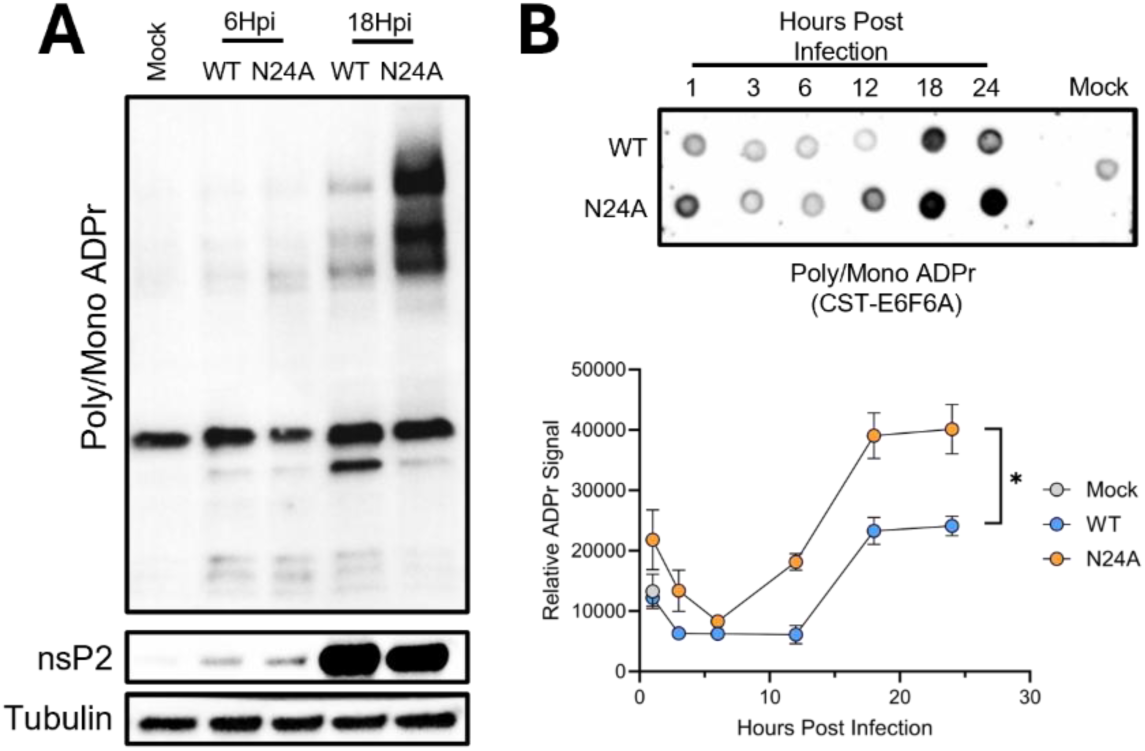
CHIKV nsP3 macrodomain activity limits RNA ribosylation in infected mammalian cells. Human fibroblasts infected with WT or N24A virus at an MOI of 3 and cell lysates were collected at 6 or 18 hours after infection. (A) The levels of viral protein expression (nsP2) and overall levels of protein ribosylation (Poly/Mono ADPr) were measured by Western blot. (B) Cells were infected as in A, and levels of RNA ribosylation in total cellular RNA was measured via dot blot (N=3, *=p<0.05).

Previous studies found that the purified CHIKV nsP3 macrodomain can remove ADPr from model RNA templates in cell-free systems (12). To determine if CHIKV nsP3 macrodomain also impacted RNA ribosylation in infected mammalian cells, human fibroblasts were infected with wild type CHIKV or the N24A mutant, and the ribosylation of total cellular RNA was measured by dot blot. Similar to protein ribosylation, the level of ADPr RNA was significantly increased in N24A infected cells as compared to wild type CHIKV infection (Figure 3B), with the largest difference occurring at later times after infection when viral RNA is the predominant RNA species in the cell. These results show that CHIKV nsP3 macrodomain activity limits the accumulation of ribosylated RNA during viral infection of mammalian cells.

### Viral RNA is Ribosylated in Mammalian Cells and in Virions

The result in Figure 3B showing higher levels of ribosylated RNA immediately after infection with N24A virus cells as compared to wild type virus, despite the presence of an equal number of intracellular viral genomes, suggested that the encapsidated N24A viral genomes themselves might be ribosylated. We therefore purified WT or N24A virions by ultracentrifugation through a sucrose density cushion, extracted RNA from the purified virus stocks, and measured ribosylated RNA levels by dot blot. N24A virion RNA was significantly more ribosylated than WT (Figure 4A, B), showing that ribosylated RNA can be packaged into CHIKV virions.

**Figure 4.**
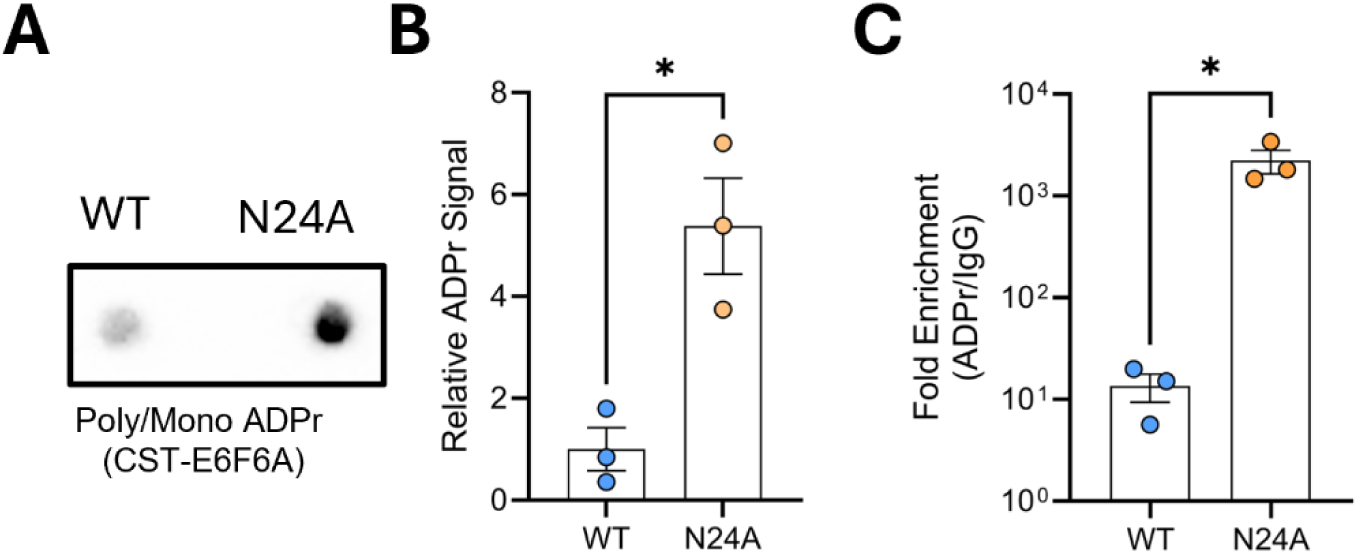
CHIKV genomes are ribosylated. (A) Virion RNA was extracted from purified CHIKV virus stocks, and levels of ribosylation were measured by dot blot. A representative is shown. (B) The results from three independent experiments described in A were quantified via densitometry. (C) Purified virion RNA from wild type (WT) or N24A virus (N24A) was incubated with a control antibody (IgG) or an antibody specific to poly/mono ADP ribose (ADPr). Antibody:RNA complexes were captured by immune precipitation, and the amount of co-purifying viral RNA was quantified by RT-qPCR. The fold enrichment of viral RNA in ADPr-specific immune complexes versus control immune complexes is shown (N=3, *=p<0.05). (N=3, *=p<0.05).

To confirm the above result we incubated purified viral RNA from WT and N24A virions with an antibody specific for ADPr or an IgG control and quantified the number of viral genomes co-purifying with each antibody by quantitative reverse transcriptase PCR (RT-qPCR). Both wild type and N24A genomes were recovered at levels above the control samples, indicating that both are ribosylated to some level. However viral genomes were significantly enriched in N24A samples incubated with the ADPr-specific antibody as compared to wild type virus, demonstrating that CHIKV RNA is ribosylated during infection and that the CHIKV macrodomain suppresses viral RNA ribosylation during infection (Figure 4C).

### PARP12 Ribosylates Viral RNA

To identify the cellular enzyme(s) that ribosylate viral RNA, we used siRNA to deplete select PARPs previously shown to regulate RNA ribosylation in mammalian cells (PARPs 10, 11, 12, 15)(12), and then measured the effect on the ribosylation of purified virion RNA and viral protein expression. Reducing the expression of PARPs 10, 11, 15 and TRPT1 had no effect on the expression of viral non-structural (nsP2) or structural (capsid) protein expression (Figure 5A). In contrast, depletion of PARP12 increased nsP2 and capsid RNA levels compared to a non-targeting siRNA control (Figures 5B, C, respectively). Analysis of virion RNA from virus stocks grown on cells depleted of each PARP alone or all in combination revealed a striking loss of virion RNA ribosylation after PARP12 depletion or the combined depletion of all PARPs together (Figure 5D, E). These data support the conclusion PARP12 is necessary for efficient ribosylation of viral RNA in mammalian cells.

**Figure 5.**
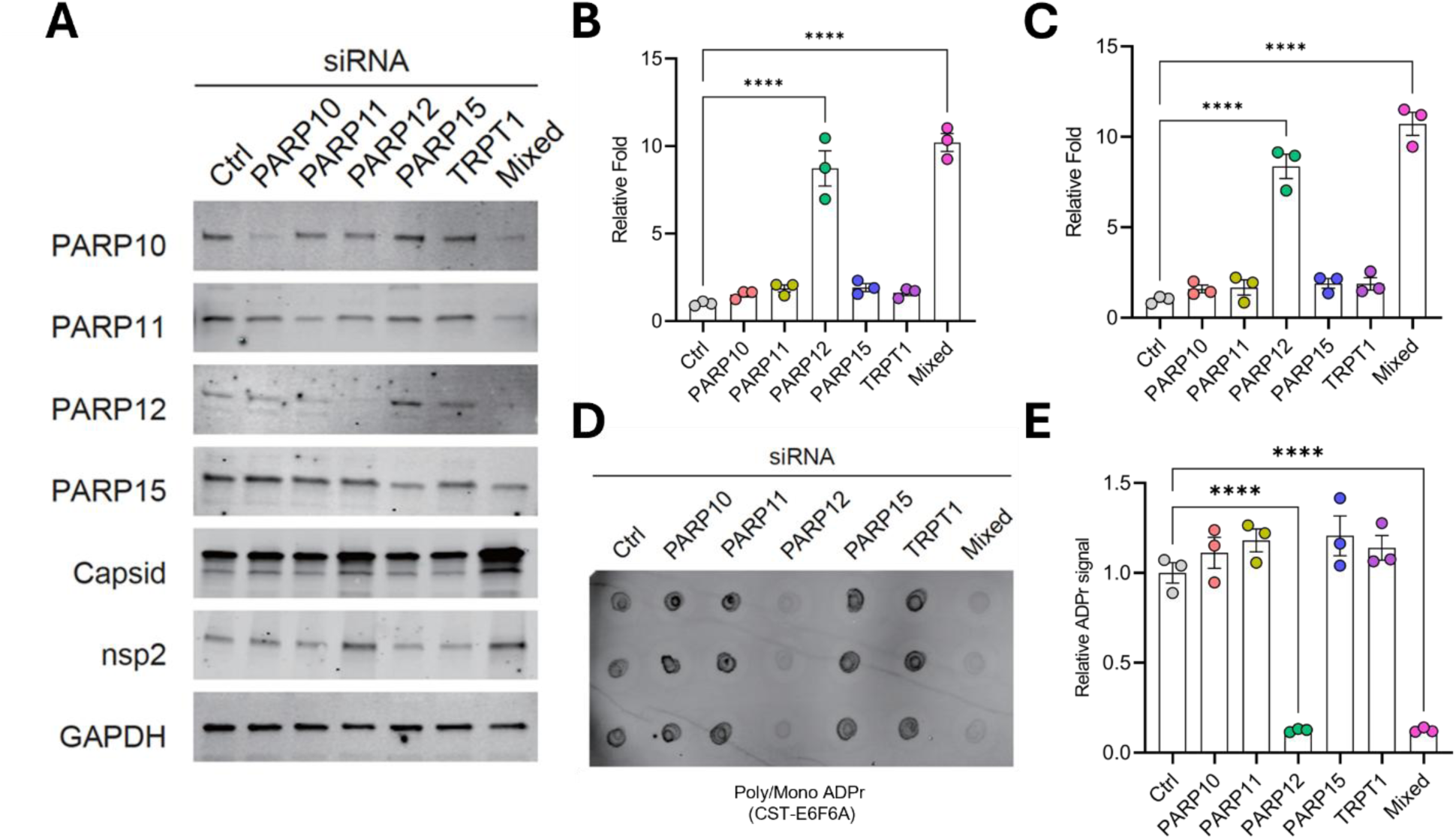
PARP12 ribosylates viral RNA in mammalian cells. (A) Human fibroblasts were transfected with a non-targeting siRNA (Ctrl), siRNA specific to the indicated PARPs, or a pooled library of all siRNAs (Mixed). Cells were infected with N24A virus at an MOI of 3 and collected at 24 hours post infection. The expression of the indicated proteins was measured by Western blot. (B) The abundance of viral RNA in siRNA treated cells was measured by RT-qPCR using primers specific for the CHIKV nsP2 non-structural protein or (C) the capsid structural gene. (D) The ribosylation levels of virion RNA from siRNA treated cells was measured by dot blot and (E) quantified by densitometry (N=3, ****=p<0.0001).

### Ribosylation Decreases the Translation Efficiency of Viral RNA

In cell-free systems, RNA can be ribosylated on its 5’ end in place of the 7-methylguanosine mRNA cap, inhibiting RNA translation (12, 30). To measure the effect of viral RNA ribosylation on viral protein synthesis, we isolated RNA from wild type or N24A virions and measured the ability of each to be translated in rabbit reticulocytes lysates. In this experiment we used a CHIKV reporter virus encoding a nanoluciferase (nLuc) reporter gene as part of the non-structural open reading frame (31). nLuc activity in this assay thus serves as a direct measure of viral RNA translation. As controls we also included in vitro transcribed, capped full length wild type and N24A genomes. Wild type virion RNA and in vitro transcribed wild type and N24A full length genomes were translated at similar rates, demonstrating that the presence of the N24A mutation alone did not affect protein synthesis. However the translation of N24A virion RNA was significantly reduced (Figure 6A), demonstrating that the defect in the translation of N24A virion RNA was due to the presence of ADPr rather than the result of a non-specific defect. To determine if ribosylation also inhibited CHIKV RNA translation in the context of infection, human fibroblasts were infected with N24A or wild type CHIKV expressing nLuc in the presence of SR-42718 (32), a potent and specific small molecule inhibitor of the CHIKV RNA-dependent RNA polymerase (RdRP) or cycloheximide (CHX), an inhibitor of global translation, and then measured nLuc expression over the first five hours after infection. In the presence of SR-42718, nLuc expression is derived solely from incoming viral genomes as de novo viral RNA synthesis is inhibited. SR-42718 decreased nLuc expression after wild type infection, consistent with inhibition of the RdRP. In control cells, nLuc expression was significantly lower in N24A infected fibroblasts as compared to cells infected with wild type virus, and the same was true in the presence of SR-42718 (Figure 6B), indicating that the input N24A genomic RNA is translated less efficiently than wild type genomic RNA. Together with the results from the in vitro translation studies, these results show that ribosylation of the viral genome inhibits its translation in mammalian systems.

**Figure 6.**
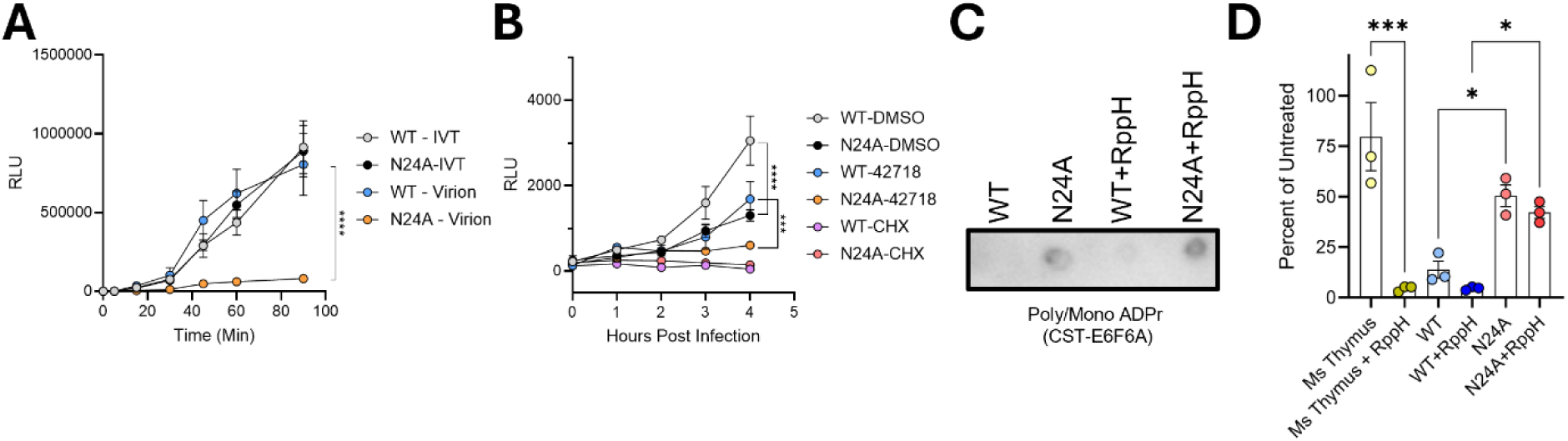
Ribosylation decreases the translation efficiency of CHIKV RNA. (A) Capped, in vitro transcribed (IVT) genomic CHIKV RNA or purified virion RNA (Virion) from wild type (WT) or N24A mutants (N24A) virus stocks expressing a nanoluciferase reporter gene were incubated with nuclease free rabbit reticulocyte lysates for the indicated times, at which nanoluciferase activity was quantified to measure the translation of each viral genome (n=3). (B) Human fibroblasts were infected with an equivalent number of viral genomes of WT or N24A nanoluciferase reporter viruses as in A and treated with cycloheximide (CHX; 100 µg/mL) or SRI-42718 (50 µM) where indicated (n=3). (C) Purified virion RNA from WT and N24A virus stocks was treated with buffer or RppH (10U) for 1 hour. Ribosylated RNA levels were measured by dot blot. (D) Mouse thymus RNA (Ms thymus) or wild type (WT) or N24A virion RNA was treated with buffer or RppH (10U) for 1 hour followed by 2U of XRN1 for 1 hour. Viral genome abundance was quantified by RT-qPCR. The graph shows the percent of input RNA remaining after XRN1 digestion. The results of a representative experiment are shown in panels C and D (*=p<0.05, ***=p<0.001,).

Previous cell-free experiments found that RNA can be ribosylated on a free 5’ monophosphate in place of the 5′ 7-methylguanosine cap, and 5’ end ribosylation significantly decreases translation and protects RNA from degradation by 5’ to 3’ exoribonucleases such as XRN1 (12). To determine if viral RNA was ribosylated on the 5’ end, we treated virion RNA with the decapping enzyme RppH followed by digestion with XRN1 and measured the amount of viral RNA remaining after each step by RT-qPCR. Viral RNA was ribosylated at similar levels in RppH and buffer treatment, demonstrating that RppH does not have ribosylhydrolase activity (Figure 6C). As expected, the control, capped RNA was minimally susceptible to degradation by XRN1 in the absence of prior RppH treatment. However the vast majority of the control RNA was degraded by XRN1 after prior incubation with RppH, indicating the control RNA has a 5’ m7G cap and removing the cap with RppH treatment makes the RNA susceptible to degradation by XRN1. We found that approx. 80-90% of wild type virion RNA was degraded by XRN1 in the absence of RppH pre-treatment, consistent with prior studies showing that the majority of wild type virion RNA is uncapped (33). Prior incubation of wild type virion RNA with RppH led to a further reduction in the levels of wild type RNA remaining, consistent with the XRN1 resistant RNA having a 5’ m^7^G cap. In contrast, significantly more N24A virion RNA was resistant to XRN1 degradation, whether or not the RNA was previously incubated with RppH (Figure 6D). These data show N24A viral RNA is protected from XRN1 degradation in a cap-independent manner, and is thus likely ribosylated on the 5’ end. Together these data show that ribosylation inhibits the translation of viral RNA in cell-free systems and in infected human fibroblasts and that the defect in translation is likely caused by ADP ribosylation at the 5’end of the RNA.

### RNA Ribosylation Decreases RNA Stability During Infection

As RNA modifications can impact RNA stability, we next determined if RNA ribosylation impacts RNA degradation in mammalian cells by measuring the decay of CHIKV RNA after infection with wild type virus or the N24A mutant. Virus was allowed to adsorb and enter infected cells for one hour, and the protein synthesis inhibitor cycloheximide or the viral RdRP inhibitor SR-42718 was then added to the cultures to prevent de novo RNA synthesis. The abundance of viral RNA as a function of time was then measured by RT-qPCR. Consistent with previous studies (33), there was an initial rapid decay of wild type viral RNA over the first hour after infection, which continued, albeit more slowly, as time progressed (Figure 7A). Cycloheximide treatment slowed viral RNA decay to a similar extent after infection with either wild type or N24A virus. However viral RNA levels decreased significantly more rapidly after infection with the N24A virus as compared to wild type in SR-42718 treated cells (Figure 7A, B). These data suggest that ribosylation enhances viral RNA decay independent of its effects on translation or viral genome replication.

**Figure 7.**
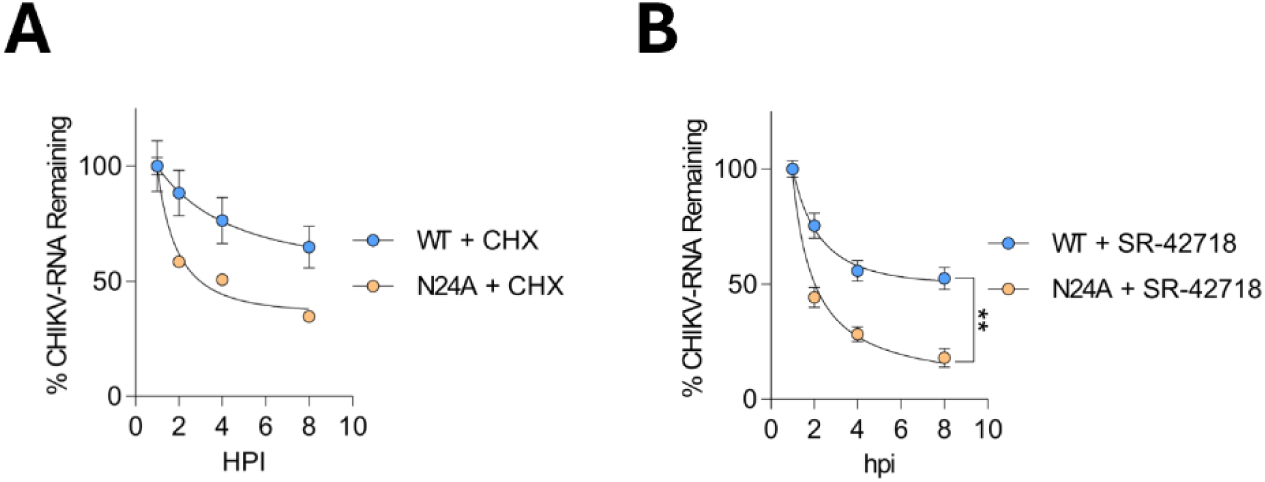
Viral RNA ribosylation enhances RNA decay. (A, B) Human fibroblasts were infected with an equal number of genomes of wild type (WT) or the N24A mutant virus. Cells were treated with cycloheximide (CHX; 100µg/mL) or SR-42718 (50 µM) to prevent de novo viral RNA synthesis. Viral genome abundance was quantified by RT-PCR at the indicated times after infection. The amount of input RNA remaining at each as a percent of viral genome abundance at 1 hour after infection is shown (N=3, **=p<0.01).

### RNA Ribosylation Increases Antiviral Gene Expression in Response to Infection

Beyond direct effects on RNA stability and translation efficiency, RNA modifications can also be recognized by cells as danger- or pathogen-associated molecular patterns (DAMPS and PAMPs, respectively), which trigger the expression of genes that function to resolve cell stress. In viral infections, PAMP recognition leads to increased expression of antiviral effector genes such as interferon stimulated genes (ISGs) that control pathogen replication. To determine if ribosylated RNA acts as a PAMP, we measured the expression of exemplar ISGs after infection of human fibroblasts with N24A or wild type CHIKV. Two CHIKV replication inhibitors, SR-42718 (32) or the alphavirus nsP2 protease inhibitor RA-2034 (34), were included in some samples to allow us to specifically measure changes in ISG induction triggered by incoming, rather than newly replicated, viral genomes. N24A viral genomes accumulated to lower levels than wild type viral genomes in the absence of the inhibitors, consistent with the defect in virus replication for the N24A virus (Figure8A, Figure 2). Both replication inhibitors prevented new viral genome accumulation in cells infected with either virus, as expected (Figure 8A). Significantly higher ISG levels were observed in cells infected with N24A virus as compared to wild type virus, and similar results were observed in the presence of the CHIKV replication inhibitors (Figure 8B-E). These results indicate that CHIKV macrodomain activity is required to suppress induction of the antiviral response and suggest that ribosylated viral RNA is a novel PAMP sensed by infected cells to initiate an antiviral response.

**Figure 8.**
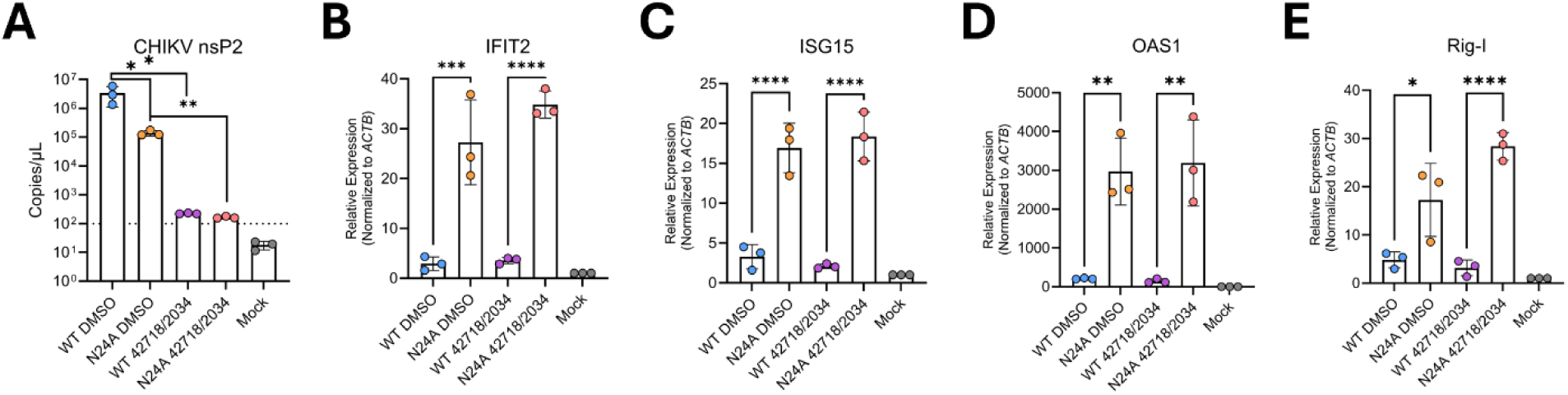
Ribosylated viral RNA triggers an enhanced cellular antiviral response. (A) Human fibroblasts were infected with an equal number of wild type (WT) or N24A mutant (N24A) viral genomes as in figure 7. Infected cells were treated with a viral replicase inhibitor (SR-42718; 50µM) in combination with an inhibitor of the viral protease (RA-2034; 50 µM), or vehicle alone (DMSO). Cells were collected at 12 hours after infection, and the amount of intracellular viral RNA was quantified by RT-qPCR using the absolute quantification method. (B-E) The fold increase in the expression of the indicated genes was measured relative to the mock sample using the ΔΔCt method normalized to actin expression (N=3, *=p<0.05, **=p<0.01, ***=p<0.001, ****=p<0.0001).

## DISCUSSION

Previous studies show that cell stress induces RNA ribosylation in mammalian cells, and cell-free studies have identified potential impacts of ribosylation in regulating RNA metabolism and translation. Here we demonstrate, to our knowledge, the first functional roles for RNA ribosylation in mammalian cells. We find that RNA ribosylation serves as a novel cellular defense during viral infection by inhibiting viral RNA translation, inducing viral RNA decay, and stimulating the innate antiviral response to infection. Our data provide new insights into how RNA ribosylation integrates into known cellular antiviral pathways suggesting a central role in the response to infection with multiple viruses. More broadly, our results provide a framework for investigating how RNA ribosylation may more generally regulate cell biology during periods of cell stress.

One key finding from our studies is that ribosylated viral RNA is poorly translated, both in cell-free systems and in human fibroblasts. Our data suggest that ADPr is added to the 5’ terminal end of viral genomes in place of the 5’ mRNA cap. These results are consistent with those from cell-free systems where the predominant ADPr modification of RNA by human PARPs occurs on 5’ terminal monophosphates of uncapped RNAs. Together with the known requirement for the 5’ mRNA cap in the translation of incoming CHIKV genomes (35), our data suggest a model in which the presence of ADPr in place of the 5’ mRNA cap reduces the recruitment of translation initiation factors needed for viral protein synthesis. We hypothesize that ribosylated viral RNA fails to recruit the cellular cap binding protein eIF4E, which is needed for the formation of translation initiation complexes that recruit ribosomes to viral RNAs. In the absence of translation initiation complex recruitment, ribosylated viral RNAs are poorly translated, limiting the expression of viral replicase proteins and decreasing virus replication. However, further studies to define the impact of viral RNA ribosylation on translation factor recruitment will be needed to more directly test this hypothesis. While our data support the conclusion that ADPr modification occurs on the 5’ end of the viral genomes, they do not rule out a potential role for ADPr modification at additional nucleotides throughout the viral genome that could potentially limit additional steps in translation such as ribosome processivity. Biophysical studies to comprehensively identify sites of ribosylation across the viral genomes and their role in viral RNA translation will be needed to determine if RNA ribosylation may play additional roles in inhibiting protein synthesis.

Our data show that RNA ribosylation decreases RNA stability in mammalian cells, as ribosylated N24A virion RNA is more rapidly degraded than wild type viral RNA. While decreased translation efficiency often correlates with increased RNA decay, our results suggest the two phenotypes are separable, as the decay of the more ribosylated N24A was accelerated compared to non-ribosylated wild type virion RNA in the presence of the translation inhibitor cycloheximide. The mechanism underlying the accelerated decay of ribosylated N24A is currently unclear. While speculative, one potential explanation is that ribosylated RNA is specifically recognized by a cellular factor that recruits the RNA degradation machinery. Such a protein would presumably have both RNA and ADPr binding activities and interact with cellular nucleases that degrade RNA. Based on our results, future efforts to define proteins that specifically interact with ribosylated RNA and regulate RNA decay in the context of infection will likely reveal key factors in the host antiviral response.

Our data also indicate that ribosylated viral RNA acts as a PAMP, as ISG induction is significantly enhanced after infection with the N24A macrodomain mutant as compared to wild type CHIKV. As with RNA decay, the enhanced ISG response was independent of de novo viral RNA synthesis, suggesting that the incoming ribosylated RNA is the trigger for enhanced ISG expression. As above, while the identity of a ribosylated RNA sensor is unknown, we speculate that this sensor would specifically recognize and bind ribosylated RNA. When bound to ligand, PAMP sensors most often activate a common set of signaling pathways (e.g. TBK1, IKKβ kinases, etc.), which in turn activate transcription factors that stimulate ISG expression. As the ISGs preferentially induced by ribosylated N24A virion RNA are also induced by other PAMP sensors (36, 37), we expect that a sensor of ribosylated RNA would activate similar cell signaling pathways, though further studies will be needed to test this hypothesis. As before, a complete understanding of how ribosylated RNA stimulates an enhanced ISG response awaits the discovery of cellular factors that recognize ribosylated RNA.

Beyond alphavirus infection, we expect our results to have implications for additional human disease states. Multiple human viral pathogens encode functional macrodomains, including multiple coronaviruses, togaviruses (38), Hepatitis E virus (39) and rubella (40). Our results suggest RNA ribosylation could play similar roles in limiting their replication. Interestingly, the SARS-CoV-2 nsP3 protein contains a macrodomain and RNA binding domain, similar to the alphavirus nsP3 protein. In cell-free systems the SARS-CoV nsP3 macrodomain also hydrolyze ADPr from RNA (13) and SARS-CoV-2 nsP3 mutants induce elevated antiviral gene expression (13). Perhaps RNA ribosylation generally triggers an antiviral response that limits virus replication in mammalian cells, similar to prokaryotes where RNA ribosylation plays a key role defending against bacteriophage infection (11, 41).

Outside the context of viral infection, RNA ribosylation is induced in response to a variety of cell stressors in mammalian cells, suggesting it may play similar roles in other disease states. ADP ribosylation is implicated in a wide range of human diseases including cardiovascular, inflammatory, and autoimmune disease, neurodegenerative disorders, and cancer. While to date the ribosylation of proteins and DNA has been viewed as the driving event in these disease states, our results suggest dysregulated RNA ribosylation may also contribute to disease, potentially providing additional avenues for therapeutic intervention.

## DATA AVAILABILITY

All relevant data is included in the manuscript.

## SUPPLEMENTARY DATA

**Supplemental Figure 1.**
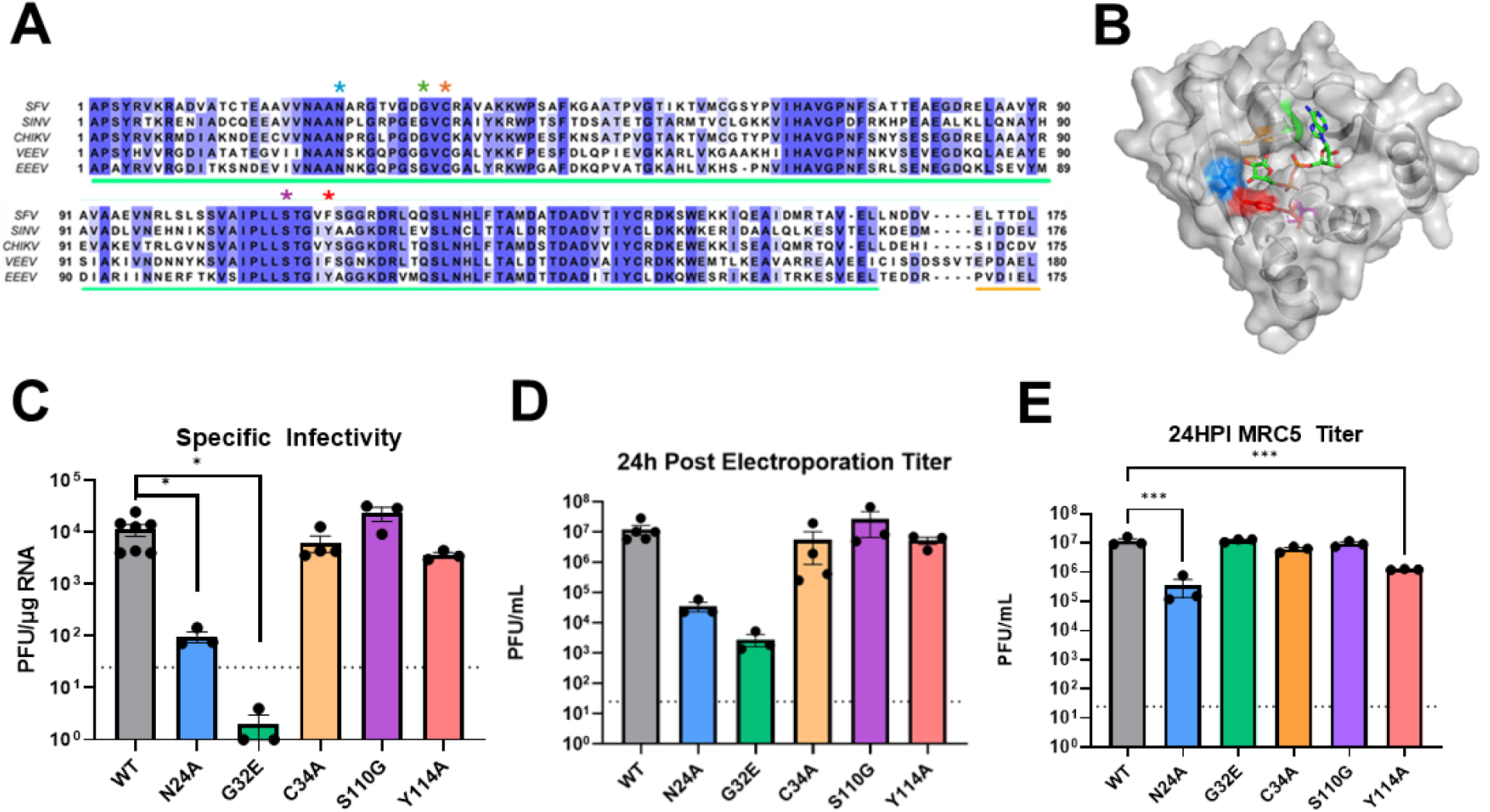
Mutagenesis of nsP3 macrodomain in chikungunya virus. (A) Alignment of amino acids of the macrodomain of the indicated alphaviruses. Mutations tested in C-D are indicated with colored asterisks (blue = N24A, green = G32E, orange = C34A, purple = S110G, red = Y114A). (B) Alphafold3 structural model of the CHIKV macrodomain with ADP-ribose. Shading corresponds to the mutation sites indicated in A. (C) The impact of the indicated mutations to the CHIKV nsP3 macrodomain on viral specific infectivity. (D) The amount of cell free infectious virus at 24 hours after electroporation was by plaque assay. (E) Human fibroblasts were infected with the indicated mutants (MOI=0.1) and viral titers in culture supernatants were quantified at 24 hours after infection by plaque assay (N=3, *=p<0.05, ***=p<0.001).

## ACKNOWLEDGEMENTS

We wish to thank members of the Moorman and Heise labs for helpful discussions. We also wish to thank Dr. Ralph Baric, Dr. Tim Sheahan and Dr. Stan Lemon for helpful conversations. We would like to thank Dr. Dan Streblow and Dr. Richard Whitley for kindly sharing reagents.

## REFERENCES

1. Boo, S.H. and Kim, Y.K. (2020) The emerging role of RNA modifications in the regulation of mRNA stability. Exp. Mol. Med., 52, 400–408.

2. Delaunay, S., Helm, M. and Frye, M. (2024) RNA modifications in physiology and disease: towards clinical applications. Nat. Rev. Genet., 25, 104–122.

3. Cui, L., Ma, R., Cai, J., Guo, C., Chen, Z., Yao, L., Wang, Y., Fan, R., Wang, X. and Shi, Y. (2022) RNA modifications: importance in immune cell biology and related diseases. Signal Transduct. Target. Ther., 7, 334.

4. Roundtree, I.A., Evans, M.E., Pan, T. and He, C. (2017) Dynamic RNA modifications in gene expression regulation. Cell, 169, 1187–1200.

5. Tong, J., Zhang, W., Chen, Y., Yuan, Q., Qin, N.-N. and Qu, G. (2022) The emerging role of RNA modifications in the regulation of antiviral innate immunity. Front. Microbiol., 13, 845625.

6. Thompson, M.G., Sacco, M.T. and Horner, S.M. (2021) How RNA modifications regulate the antiviral response. Immunol. Rev., 304, 169–180.

7. Suskiewicz, M.J., Prokhorova, E., Rack, J.G.M. and Ahel, I. (2023) ADP-ribosylation from molecular mechanisms to therapeutic implications. Cell, 186, 4475–4495.

8. Krieg, S., Pott, F., Potthoff, L., Verheirstraeten, M., Bütepage, M., Golzmann, A., Lippok, B., Goffinet, C., Lüscher, B. and Korn, P. (2023) Mono-ADP-ribosylation by PARP10 inhibits Chikungunya virus nsP2 proteolytic activity and viral replication. Cell. Mol. Life Sci., 80, 72.

9. Ryan, A.P., Delgado-Rodriguez, S.E. and Daugherty, M.D. (2025) Zinc-finger PARP proteins ADP-ribosylate alphaviral proteins and are required for interferon-γ-mediated antiviral immunity. Sci. Adv., 11, eadm6812.

10. Du, Q., Miao, Y., He, W. and Zheng, H. (2023) ADP-Ribosylation in Antiviral Innate Immune Response. Pathogens, 12.

11. Vassallo, C.N., Doering, C.R. and Laub, M.T. (2024) Anti-viral defence by an mRNA ADP-ribosyltransferase that blocks translation. Nature, 636, 190–197.

12. Weixler, L., Feijs, K.L.H. and Zaja, R. (2022) ADP-ribosylation of RNA in mammalian cells is mediated by TRPT1 and multiple PARPs. Nucleic Acids Res., 50, 9426–9441.

13. Munnur, D., Bartlett, E., Mikolčević, P., Kirby, I.T., Rack, J.G.M., Mikoč, A., Cohen, M.S. and Ahel, I. (2019) Reversible ADP-ribosylation of RNA. Nucleic Acids Res., 47, 5658–5669.

14. Groslambert, J., Prokhorova, E. and Ahel, I. (2021) ADP-ribosylation of DNA and RNA. DNA Repair (Amst), 105, 103144.

15. Bullen, N.P., Sychantha, D., Thang, S.S., Culviner, P.H., Rudzite, M., Ahmad, S., Shah, V.S., Filloux, A., Prehna, G. and Whitney, J.C. (2022) An ADP-ribosyltransferase toxin kills bacterial cells by modifying structured non-coding RNAs. Mol. Cell, 82, 3484–3498.e11.

16. Lu, Y., Schuller, M., Bullen, N.P., Mikolcevic, P., Zonjic, I., Raggiaschi, R., Mikoc, A., Whitney, J.C. and Ahel, I. (2025) Discovery of reversing enzymes for RNA ADP- ribosylation reveals a possible defence module against toxic attack. Nucleic Acids Res., 53.

17. Mikolčević, P., Hloušek-Kasun, A., Ahel, I. and Mikoč, A. (2021) ADP-ribosylation systems in bacteria and viruses. Comput. Struct. Biotechnol. J., 19, 2366–2383.

18. Aravind, L., Zhang, D., de Souza, R.F., Anand, S. and Iyer, L.M. (2015) The natural history of ADP-ribosyltransferases and the ADP-ribosylation system. Curr. Top. Microbiol. Immunol., 384, 3–32.

19. Wyżewski, Z., Gradowski, M., Krysińska, M., Dudkiewicz, M. and Pawłowski, K. (2021) A novel predicted ADP-ribosyltransferase-like family conserved in eukaryotic evolution. PeerJ, 9, e11051.

20. Cihlova, B., Lu, Y., Mikoč, A., Schuller, M. and Ahel, I. (2024) Specificity of DNA ADP- Ribosylation Reversal by NADARs. Toxins (Basel), 16.

21. Weixler, L., Žaja, R., Ikenga, N.J., Siefert, J., Mohan, G., Aydin, G., Wijngaarden, S., Filippov, D.V., Lüscher, B. and Feijs-Žaja, K.L.H. (2025) Family-wide analysis of human macrodomains reveals novel activities and identifies PARG as most efficient ADPr- RNA hydrolase. Commun. Biol., 8, 453.

22. Vincent, H.A., Ziehr, B. and Moorman, N.J. (2017) Mechanism of Protein Kinase R Inhibition by Human Cytomegalovirus pTRS1. J. Virol., 91.

23. Weixler, L., Ikenga, N.J., Voorneveld, J., Aydin, G., Bolte, T.M., Momoh, J., Bütepage, M., Golzmann, A., Lüscher, B., Filippov, D.V., et al. (2023) Protein and RNA ADP-ribosylation detection is influenced by sample preparation and reagents used. Life Sci. Alliance, 6.

24. Cruz, C.C., Suthar, M.S., Montgomery, S.A., Shabman, R., Simmons, J., Johnston, R.E., Morrison, T.E. and Heise, M.T. (2010) Modulation of type I IFN induction by a virulence determinant within the alphavirus nsP1 protein. Virology, 399, 1–10.

25. Arend, K.C., Ziehr, B., Vincent, H.A. and Moorman, N.J. (2016) Multiple Transcripts Encode Full-Length Human Cytomegalovirus IE1 and IE2 Proteins during Lytic Infection. J. Virol., 90, 8855–8865.

26. Dickmander, R.J., Lenarcic, E.M., Sears, J.D., Hale, A.E. and Moorman, N.J. (2025) RNA-targeted proteomics identifies YBX1 as critical for efficient HCMV mRNA translation. Proc Natl Acad Sci USA, 122, e2421155122.

27. Abraham, R., Hauer, D., McPherson, R.L., Utt, A., Kirby, I.T., Cohen, M.S., Merits, A., Leung, A.K.L. and Griffin, D.E. (2018) ADP-ribosyl-binding and hydrolase activities of the alphavirus nsP3 macrodomain are critical for initiation of virus replication. Proc Natl Acad Sci USA, 115, E10457–E10466.

28. Abraham, R., McPherson, R.L., Dasovich, M., Badiee, M., Leung, A.K.L. and Griffin, D.E. (2020) Both ADP-Ribosyl-Binding and Hydrolase Activities of the Alphavirus nsP3 Macrodomain Affect Neurovirulence in Mice. MBio, 11.

29. Jayabalan, A.K., Adivarahan, S., Koppula, A., Abraham, R., Batish, M., Zenklusen, D., Griffin, D.E. and Leung, A.K.L. (2021) Stress granule formation, disassembly, and composition are regulated by alphavirus ADP-ribosylhydrolase activity. Proc Natl Acad Sci USA, 118.

30. Munir, A., Banerjee, A. and Shuman, S. (2018) NAD+-dependent synthesis of a 5’- phospho-ADP-ribosylated RNA/DNA cap by RNA 2’-phosphotransferase Tpt1. Nucleic Acids Res., 46, 9617–9624.

31. Ahmed, S.K., Haese, N.N., Cowan, J.T., Pathak, V., Moukha-Chafiq, O., Smith, V.J., Rodzinak, K.J., Ahmad, F., Zhang, S., Bonin, K.M., et al. (2021) Targeting chikungunya virus replication by benzoannulene inhibitors. J. Med. Chem., 64, 4762–4786.

32. WO2021203048A1 - Novel 2-pyrimidone analogs as potent antiviral agents against alphaviruses - Google Patents https://patents.google.com/patent/WO2021203048A1/en (14 July 2025, date last accessed).

33. Sokoloski, K.J., Haist, K.C., Morrison, T.E., Mukhopadhyay, S. and Hardy, R.W. (2015) Noncapped Alphavirus Genomic RNAs and Their Role during Infection. J. Virol., 89, 6080–6092.

34. Merten, E.M., Sears, J.D., Leisner, T.M., Hardy, P.B., Ghoshal, A., Hossain, M.A., Asressu, K.H., Brown, P.J., Tse, E.G., Stashko, M.A., et al. (2024) Identification of a cell-active chikungunya virus nsP2 protease inhibitor using a covalent fragment-based screening approach. Proc. Natl. Acad. Sci. USA, 121.

35. Broeckel, R., Sarkar, S., May, N.A., Totonchy, J., Kreklywich, C.N., Smith, P., Graves, L., DeFilippis, V.R., Heise, M.T., Morrison, T.E., et al. (2019) Src Family Kinase Inhibitors Block Translation of Alphavirus Subgenomic mRNAs. Antimicrob. Agents Chemother., 63.

36. Barral, P.M., Sarkar, D., Su, Z., Barber, G.N., DeSalle, R., Racaniello, V.R. and Fisher, P.B. (2009) Functions of the cytoplasmic RNA sensors RIG-I and MDA-5: key regulators of innate immunity. Pharmacol. Ther., 124, 219–234.

37. Loo, Y.-M., Fornek, J., Crochet, N., Bajwa, G., Perwitasari, O., Martinez-Sobrido, L., Akira, S., Gill, M.A., García-Sastre, A., Katze, M.G., et al. (2008) Distinct RIG-I and MDA5 signaling by RNA viruses in innate immunity. J. Virol., 82, 335–345.

38. Alhammad, Y.M.O. and Fehr, A.R. (2020) The Viral Macrodomain Counters Host Antiviral ADP-Ribosylation. Viruses, 12.

39. Eckei, L., Krieg, S., Bütepage, M., Lehmann, A., Gross, A., Lippok, B., Grimm, A.R., Kümmerer, B.M., Rossetti, G., Lüscher, B., et al. (2017) The conserved macrodomains of the non-structural proteins of Chikungunya virus and other pathogenic positive strand RNA viruses function as mono-ADP-ribosylhydrolases. Sci. Rep., 7, 41746.

40. Stoll, G.A., Nikolopoulos, N., Zhai, H., Zhang, L., Douse, C.H. and Modis, Y. (2024) Crystal structure and biochemical activity of the macrodomain from rubella virus p150. J. Virol., 98, e0177723.

41. Mets, T., Kurata, T., Ernits, K., Johansson, M.J.O., Craig, S.Z., Evora, G.M., Buttress, J.A., Odai, R., Wallant, K.C., Nakamoto, J.A., et al. (2024) Mechanism of phage sensing and restriction by toxin-antitoxin-chaperone systems. Cell Host Microbe, 32, 1059–1073.e8.

